# RIF1 integrates DNA repair and transcriptional requirements during the establishment of humoral immune responses

**DOI:** 10.1101/2023.07.18.549543

**Authors:** Ali Rahjouei, Eleni Kabrani, Maria Berruezo-Llacuna, Robert Altwasser, Veronica Delgado-Benito, Rushad Pavri, Michela Di Virgilio

## Abstract

The establishment of protective immune responses relies on the ability of terminally differentiated B cells to secrete a broad variety of antigen-specific antibodies with different effector functions. RIF1 is a multifunctional protein that promotes antibody isotype diversification *via* its DNA end protection activity during class switch recombination (CSR). In this study, we showed that RIF1 ablation resulted in increased plasmablast (PB) formation *ex vivo* and enhanced terminal differentiation into plasma cells (PCs) upon immunization. Mechanistically, this phenotype is independent from RIF1’s role in DNA repair and CSR, and reflects its ability to modulate the transcriptional status of a subset of BLIMP1 target genes. Therefore, in addition to promoting antibody isotype diversification, RIF1 fine-tunes the kinetics of late B cell differentiation, thus providing an additional layer of control in the establishment of humoral immunity.

## Introduction

During early B cell development in the bone marrow, common lymphoid progenitors develop into immature B cells *via* a step-wise process linked to the rearrangement of their antibody receptor/immunoglobulin (Ig) genes by V(D)J recombination ^1,2^. Immature B cells expressing a functional and non-autoreactive B cell receptor (BCR) migrate to the periphery, where they complete their maturation into quiescent resting cells ^1,2^. The encounter of these naïve mature B cells with their cognate antigen in secondary lymphoid organs triggers late B cell differentiation into effector cells, often accompanied by further BCR/Ig diversification *via* somatic hypermutation (SHM) and class switch recombination (CSR) ^3^. SHM and CSR provide the molecular bases for generating high-affinity and isotype-switched BCRs/Igs, respectively ^4^. At the cellular level, the combined effect of BCR engagement, helper T and follicular dendritic cell-derived signals induces the formation of specialized microstructures known as germinal centers (GCs), where activated B cells undergo a major proliferative burst, selection of high-affinity BCR variants, and differentiation into either antibody-secreting cells (ASCs, plasmablasts (PBs) and plasma cells (PCs)) or memory B cells ^5^. B cell activation can also lead to PB generation at extrafollicular sites, which are also characterized by B cell proliferation and represent the main source of ASCs in T-cell independent responses ^6^. BLIMP1 (encoded by the *Prdm1* gene) is the transcriptional master regulator of late B cell differentiation ^7^. BLIMP1 is dispensable for the initiation of the ASC program but is necessary for the establishment of the terminally differentiated PC phenotype, mainly through suppression of its target genes ^8–11^. The generation of PBs and long-lived PCs capable of secreting high-affinity Igs of different classes provides the foundation for the establishment of protective humoral responses.

RIF1 (Rap1-Interacting Factor 1 Homolog / Replication Timing Regulatory Factor 1) is a multifunctional protein initially identified in budding yeast as a regulator of telomere length homeostasis ^12^. In mammalian cells, RIF1 contributes to preserving genome stability during both DNA replication and repair. Under conditions of DNA replication stress, RIF1 protects nascent DNA at stalled forks from degradation, facilitating their timely restart ^13–16^. During the repair of DNA double-strand breaks (DSBs), RIF1 participates in the 53BP1-Shieldin cascade that protects the broken DNA ends against nucleolytic resection, thus influencing the choice of which DSB repair pathway to engage ^17–21^. In addition to these genome-protective functions, RIF1 plays a central role in the control of DNA replication timing programs in both yeast and higher eukaryotes ^22–25^. Several studies have also implicated RIF1 in early mouse development ^26–30^. This role appears independent from RIF1’s various activities in DNA metabolism, and is mediated by its ability to modulate the transcriptional networks responsible for embryonic stem cell state stability and differentiation ^26–30^.

Isotype diversification by CSR occurs *via* a deletional recombination reaction at the Ig heavy chain (*Igh*) locus, which replaces the constant (C) region of the IgM/IgD basal isotype with one of the downstream C regions encoding for the different classes (IgG, IgE or IgA) ^4^. The process is initiated by the B cell-specific enzyme Activation-Induced Deaminase (AID), whose activity at the *Igh* locus leads to the formation of DSBs that are obligate intermediates in the reaction ^3^. These programmed breaks need to be physiologically protected from extensive resection to enable productive repair events and CSR ^31^. Due to its ability to inhibit DSB end processing, RIF1 is required for repair of *Igh* breaks, and hence for efficient Ig isotype diversification ^17–19^. In this study, we report a novel role for RIF1 in the regulation of humoral immunity. We discovered that RIF1 expression is upregulated in mature B cells following activation, where it binds cis-regulatory elements of genes involved in B cell function and differentiation. RIF1 deficiency skews the transcriptional profile of activated B cells towards PBs and PCs, and is associated with an accelerated differentiation to ASCs both *ex* and *in vivo*. RIF1 directly binds several BLIMP1 target genes and counteracts their premature repression. Thus, RIF1 contributes an additional regulatory layer to the B cell differentiation program that is essential to establish secreted antibody diversity.

## Results

### *Rif1* expression is regulated during B cell differentiation

Given RIF1’s multiple roles during early development and in differentiated somatic cells ^26,27,29,30,32,33^, we asked whether RIF1 contributes functions beyond the regulation of DNA end processing and repair in B cells. To do so, we first monitored *Rif1* expression levels across B cell lineage developmental stages using the Immunological Genome Project (ImmGen) transcriptomics data ^34^. *Rif1* expression varied considerably in the different B cell subtypes, with GC cells exhibiting the highest levels (Fig. 1a). In contrast, the expression of RIF1’s up- and downstream partners in DNA end protection, 53BP1 and the Shieldin (MAD2L2-SHLD1-SHLD2-SHLD3) complex, respectively, did not show any major changes during B cell development and differentiation (Fig. 1a and S1a). In addition, *ex vivo* activation of isolated naïve B cells (Fig. 1b and S1b) resulted in upregulation of *Rif1* transcript levels, regardless of the stimulation condition (LPS and IL-4 (LI); LPS, BAFF and TGFβ (LBT); or LPS (L)) and resulting isotype switching (IgG1, IgG2b, or IgG3, respectively) (Fig. 1c). MAD2L2 was the only additional component of the 53BP1-RIF1-Shieldin axis to exhibit transcriptional induction after *ex vivo* stimulation (Fig. 1c and S1c). Finally, *Rif1* transcripts’ upregulation was accompanied by an increase in the expression also at the protein level, which was readily detectable at 48 h after activation, sustained at 72 h and declining thereafter (Fig. 1d). We concluded that *Rif1* expression is induced following activation of mature B cells both *in* and *ex vivo*, and that this expression dynamics does not reflect a general behavior of DSB end protection factors.

**Figure 1.**
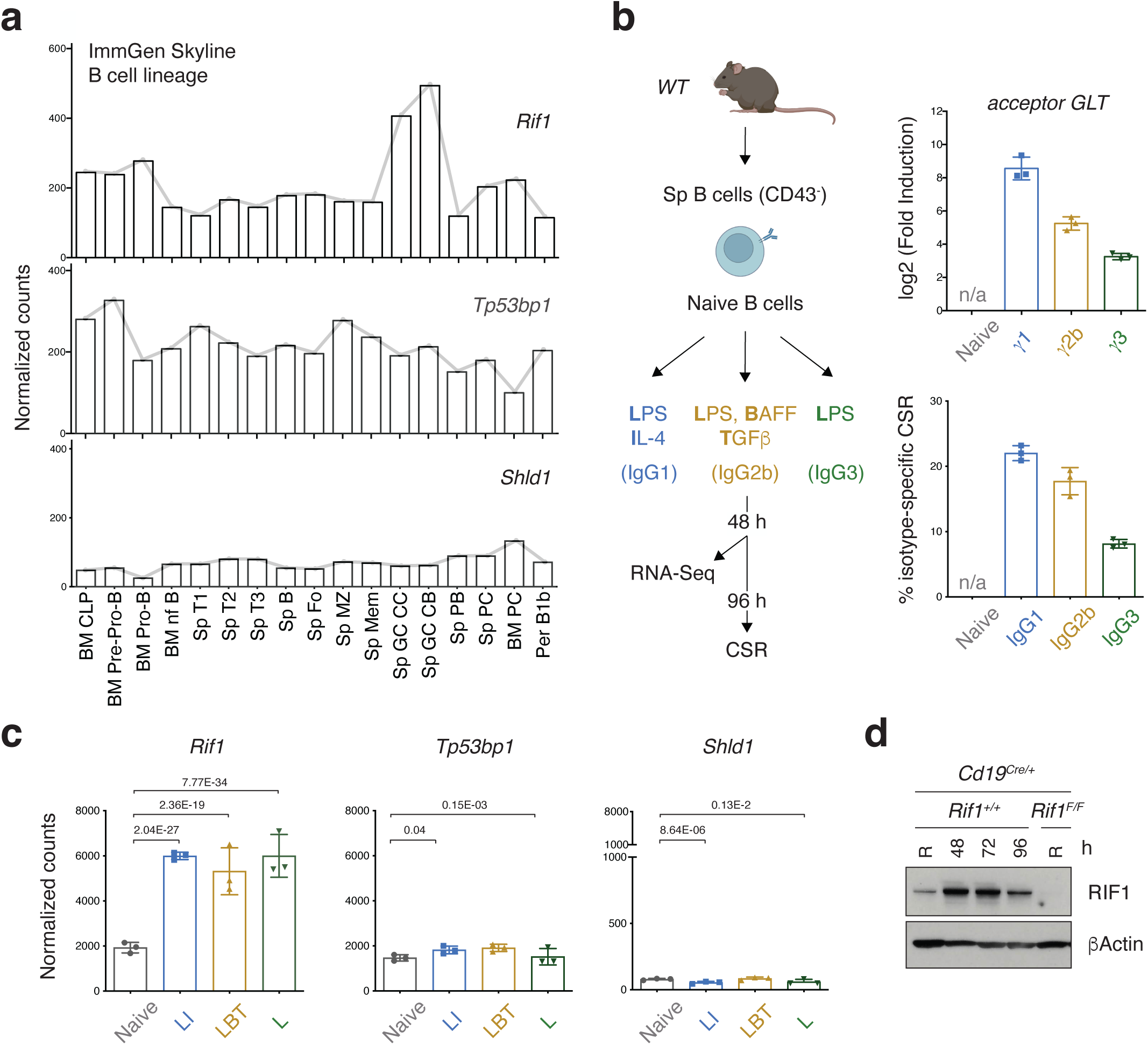
*Rif1* expression is dynamically regulated during late B cell differentiation. **a** Expression of *Rif1*, *Tp53bp1*, and *Shld1* genes across B cell lineage developmental stages as determined by the Immunological Genome Project (ImmGen) Skyline RNA-Seq analysis. BM: bone marrow; CLP: common lymphoid progenitor; nf: newly-formed; Sp: splenic; Fo: follicular; MZ: marginal zone; Mem: memory; GC: germinal center; CC: centrocytes; CB: centroblasts; PB: plasmablasts; PC: plasma cells; Per B1b: peritoneal B1b. **b** Left: Schematic representation of gene expression analysis in naïve B cells isolated from mouse spleens and stimulated *ex vivo* with LPS and IL-4 (LI cocktail), LPS, BAFF and TGFβ (LBT), or LPS only (L). Right: Levels of acceptor germline transcript (GLT) induced by LI-, LBT- or L-stimulation (expressed as fold increase over the naïve sample) and CSR efficiency to the corresponding isotype for each of the primary B cell cultures employed in the RNA-Seq analysis (n = 3 mice per stimulation condition). n/a: not applicable. **c** Expression of *Rif1*, *Tp53bp1*, and *Shld1* in naïve and LI/LBT/L-stimulated B cells. **d** Western blot analysis of *Cd19^Cre/+^* and *Rif1^F/F^Cd19^Cre/+^* purified B lymphocytes at the indicated times after activation with LI. Data is representative of two independent genotype pairs. R: Resting. Expression values in panels **a**, **b** and **c** were normalized by DESeq2. The adjusted p-value of significant differences between samples in panel **c** is indicated.

**Figure S1.**
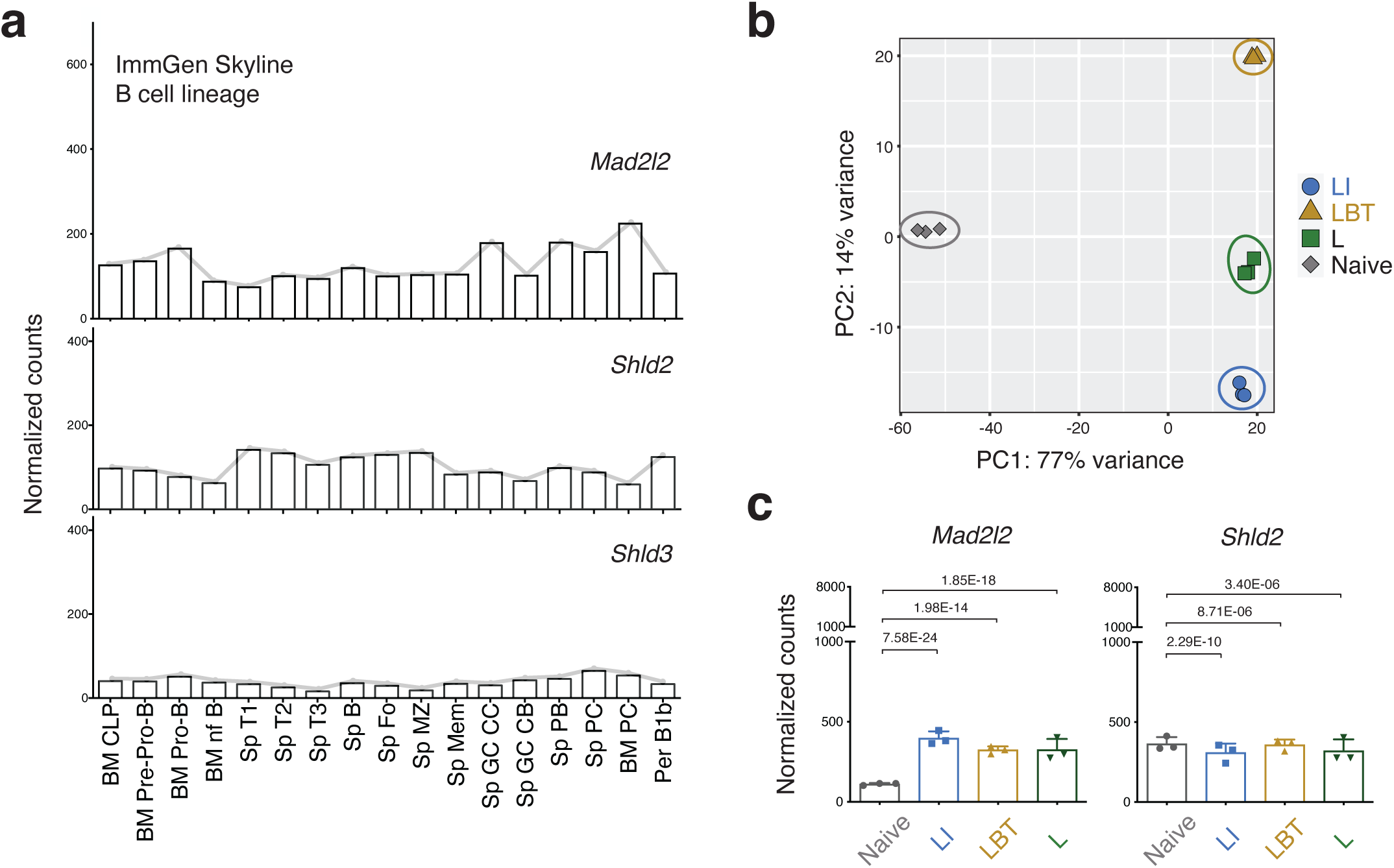
Stimulation-dependent clustering of B cell transcriptomics profiles after *ex vivo* activation. **a** Expression of *Mad2l2*, *Shld2* and *Shld3* genes across B cell lineage developmental stages as determined by the Immunological Genome Project (ImmGen) Skyline RNA-Seq analysis. BM: bone marrow; CLP: common lymphoid progenitor; nf: newly-formed; Sp: splenic; Fo: follicular; MZ: marginal zone; Mem: memory; GC: germinal center; CC: centrocytes; CB: centroblasts; PB: plasmablasts; PC: plasma cells; Per B1b: peritoneal B1b. **b** Principal component analysis for the RNA-Seq datasets of WT splenocytes before (naïve) and after 48 h stimulation with LPS and IL-4 (LI), LPS, BAFF and TGFβ (LBT), or LPS only (L). The RNA-Seq analysis was performed on three biological replicates/mice per condition. **c** Expression of *Mad2l2* and *Shld2* in naïve and LI/LBT/L-stimulated B cells. *Shld3* expression levels were below detection in all datasets. Expression values in panels **a** and **c** were normalized by DESeq2. The adjusted p-value of significant differences between samples in panel **c** is indicated. Related to Figure 1.

### RIF1 limits the *ex vivo* differentiation of activated B cells into PBs and PC-like cells

The specific upregulation of RIF1 expression in GC and *ex vivo* stimulated splenocytes (Fig. 1) suggests that RIF1’s functional repertoire in activated B cells might extend beyond its established role in the regulation of DNA end processing. In addition to SHM and CSR, activation of mature B cells triggers the late stages of B cell development ^5,6,35^. Therefore, we next investigated the potential involvement of RIF1 during ASC generation.

Stimulation of naïve B cells with LPS and IL-4 induces their *ex vivo* differentiation into PBs (evidenced by the surface expression of CD138). To assess the contribution of RIF1 to this process, we employed B lymphocyte cultures from *Rif1^F/F^Cd19^Cre/+^*mice (Fig. 1d), which conditionally ablate *Rif1* expression at the immature B cell stage ^18^. We found an over two-fold increase in the percentage of CD138^+^ cells in *Rif1^F/F^Cd19^Cre/+^*samples compared to *Cd19^Cre^* (Fig. 2a), despite similar proliferation rates between the two genotypes (Fig. 2b and ^18^). Furthermore, whereas controls exhibited a steep rise in PB formation mainly at the last cell division, RIF1-deficient cultures showed higher and increasing levels of PBs from earlier division rounds (Fig. 2b). The observed phenotype is not linked to RIF1’s role in the regulation of DNA end processing nor to the CSR defect associated to its ablation since B cell cultures from mice lacking either a functional Shieldin complex (*Shld1^-/-^*) or AID (*Aicda^-/-^*) differentiated to the same extent as controls (Fig. 2c).

**Figure 2.**
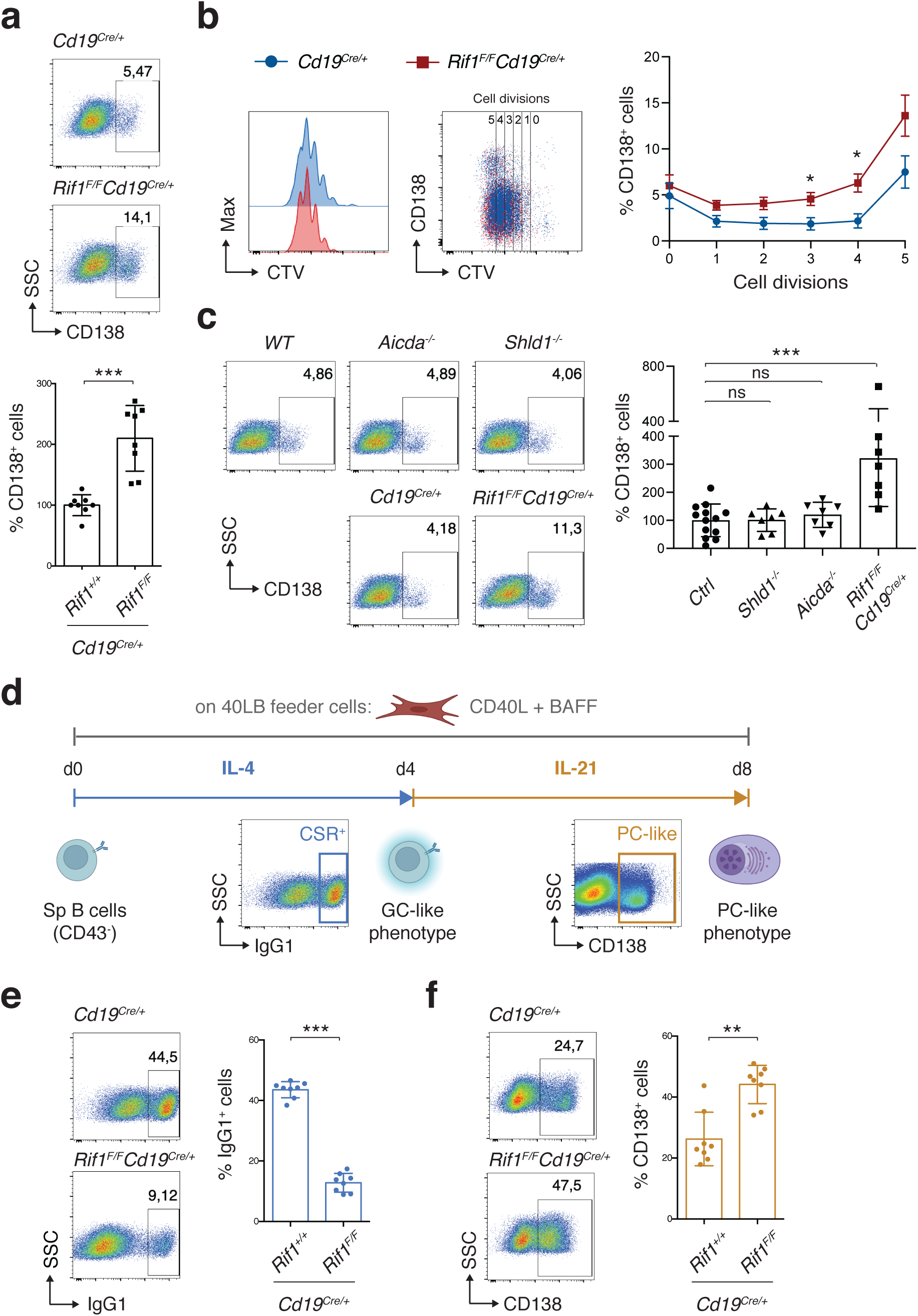
RIF1 limits the *ex vivo* differentiation of activated B cells to PBs and PC-like cells independently from its role in DNA end protection and CSR. **a** Top: Representative flow cytometry plots measuring percentage of plasmablasts (CD138+) in purified splenic B cell cultures of the indicated genotypes 96 h after activation with LPS and IL-4 (LI). Bottom: Summary graph for five independent experiments (n = eight mice per genotype), with % of CD138+ cells within each experiment normalized to the average of control mice (*Cd19^Cre/+^*), which was set to 100%. **b** Left: Representative flow cytometry plots measuring proliferation by CellTrace Violet (CTV) dilution and percentage of CD138+ cells per cell division in primary cultures of *Cd19^Cre/+^* and *Rif1^F/F^Cd19^Cre/+^* B lymphocytes stimulated with LI for 96 h. Right: Summary graph for three independent experiments (n = 6 mice per genotype). **c** Left: Representative flow cytometry plots measuring percentage of CD138+ cells in purified B cell cultures of the indicated genotypes 96 h after activation with LI. Right: Summary graph for four independent experiments (n = seven mice per genotype) with % of CD138+ cells within each experiment normalized to the average of control mice (*WT* and *Cd19^Cre/+^*: *Ctrl*), which was set to 100%. **d** Schematic representation of the iGB (induced germinal center (GC) phenotype B) cell culture system for the *ex vivo* differentiation of splenic naïve B cells to GC- and plasma cell-like cells. CD40L: CD40 ligand; d: day; Sp: splenic; CSR+: class switched cells; PC: plasma cell. **e**-**f** Left: Representative flow cytometry plots measuring CSR to IgG1 (**e**) and percentage of plasmablasts (CD138+) (**f**) in purified B cell cultures of the indicated genotypes at day 4 (**e**) and 8 (**f**), respectively, of the iGB cell culture system shown in panel **d**. Right: Summary graphs for five independent experiments (n = eight mice per genotype). Significance in panels **a**, **b**, **c**, **e** and **f** was calculated with the Mann–Whitney U test, with error bars representing SD in panels **a**, **c**, **e** and **f**, and SEM in panel **b**. Only significant differences between groups were indicated in panel **b**. ns: not significant; * = p ≤ 0.05; ** = p ≤ 0.01; *** = p ≤ 0.001.

**Figure S2.**
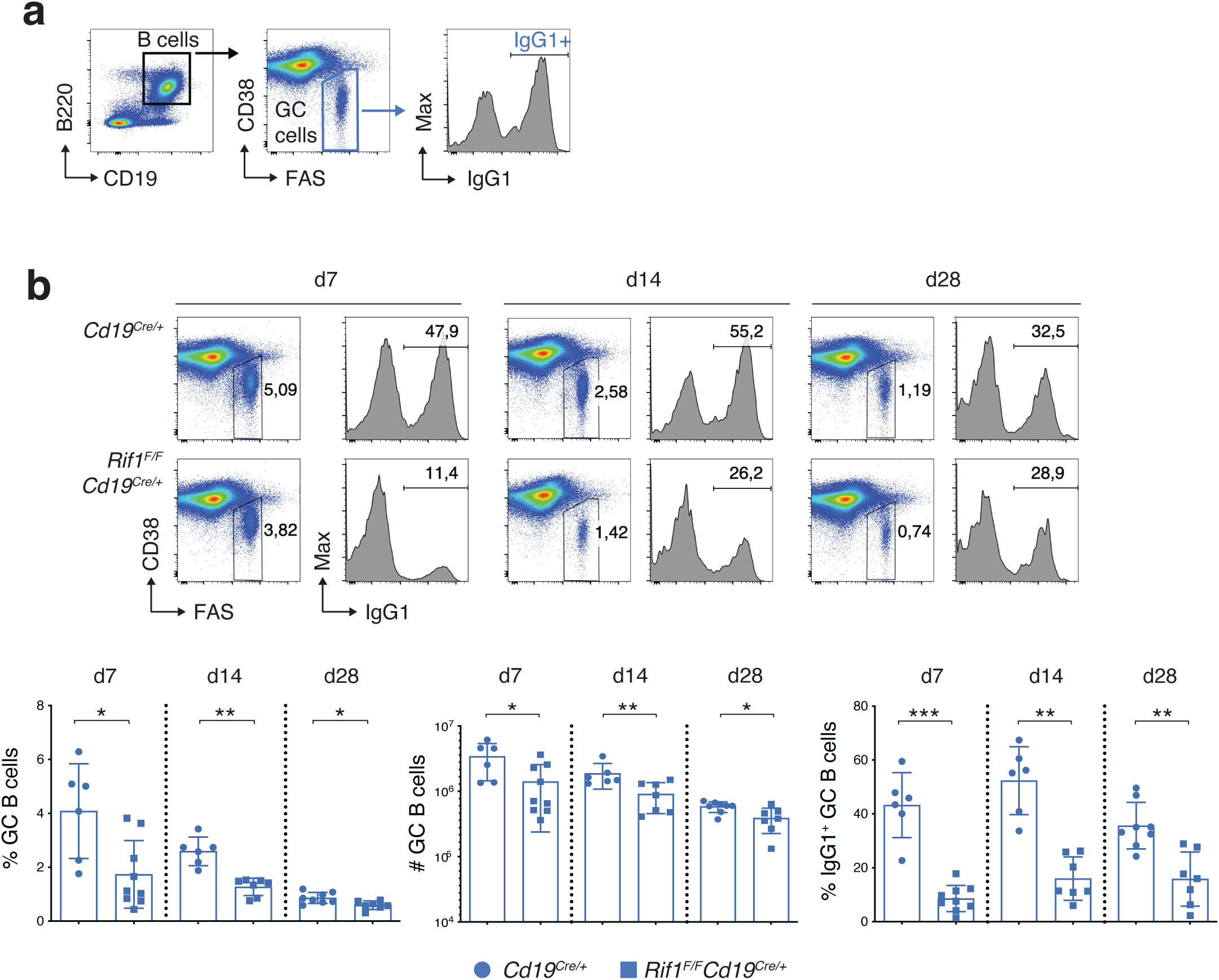
RIF1 deficiency results in a reduced number of class-switched GC B cells upon immunization. **a** Gating strategy for Germinal Center (GC) analysis. **b** Top: Representative flow cytometry plots measuring the percentage of GC cells and IgG1-class switched GC cells in spleens of *Cd19^Cre/+^* and *Rif1^F/F^Cd19^Cre/+^* mice at the indicated days after immunization. Bottom: Summary graphs for at least six mice per genotype and time point in at least three independent experiments. Significance in panel **b** was calculated with the Mann–Whitney U test and error bars represent SD. * = p ≤ 0.05; ** = p ≤ 0.01; *** = p ≤ 0.001. Related to Figure 4.

Currently, no *in vitro* setting can fully recapitulate the complexity of PC differentiation and function. However, the induced GC B (iGB) culture system ideated by Nojima *et al.* mimics the T cell-dependent generation of GC B cells and enables the *in vitro* manipulation of their fates into either memory- or long-lived PC-like cells ^36^ (Fig. 2d). We took advantage of this system to assess the contribution of RIF1 to the terminal differentiation of activated B cells *ex vivo*. At the GC-like phenotype stage (four days stimulation with IL-4 on CD40L- and BAFF-expressing feeder cells), RIF1 deficiency resulted in the expected severe defect in CSR (Fig. 2, d and e, and ^17–19)^. However, and in agreement with their increased potential to differentiate into PBs (Fig. 2, a to c), *Rif1^F/F^Cd19^Cre^* B cells showed a near two-fold increase in the percentage of PC-like cells after prolonged culturing in the presence of IL-21 (Fig. 2, d and f).

Altogether, these data indicate that RIF1 deficiency accelerates the *ex vivo* differentiation of activated B cells to PBs and PC-like cells, and that this function is independent from RIF1’s role in DNA end protection and CSR.

### RIF1 deficiency does not affect GC B cell dynamics

We next asked whether RIF1 is able to modulate late B cell differentiation also *in vivo*. To this end, we first assessed the consequences of RIF1 ablation on GC physiology by comparing GC B cell dynamics in Peyer’s patches (PPs) of *Cd19^Cre/+^* and *Rif1^F/F^Cd19^Cre/+^*mice. GCs are anatomically separated into the dark zone (DZ) and light zone (LZ) ^5,6^, and cycling of positively selected GC B cells between LZ and DZ ensures further affinity maturation and clonal expansion ^5,6^. PPs are specialized secondary lymphoid structures that line the wall of the small intestine and are crucial for mucosal IgA antibody responses ^37^. PPs allow for the assessment of chronic GC activity in response to a variety of food- and microbiome-derived antigens.

Control and *Rif1^F/F^Cd19^Cre/+^* mice showed no significant difference in the percentage of PP GC B cells nor in their distribution in the LZ *versus* DZ (Fig. 3, a and b). In contrast, mice knock-out for AID exhibited the expected massive increase in GC B cells and a cellular distribution heavily skewed towards the LZ (Fig. 3b and ^38–40^).

**Figure 3.**
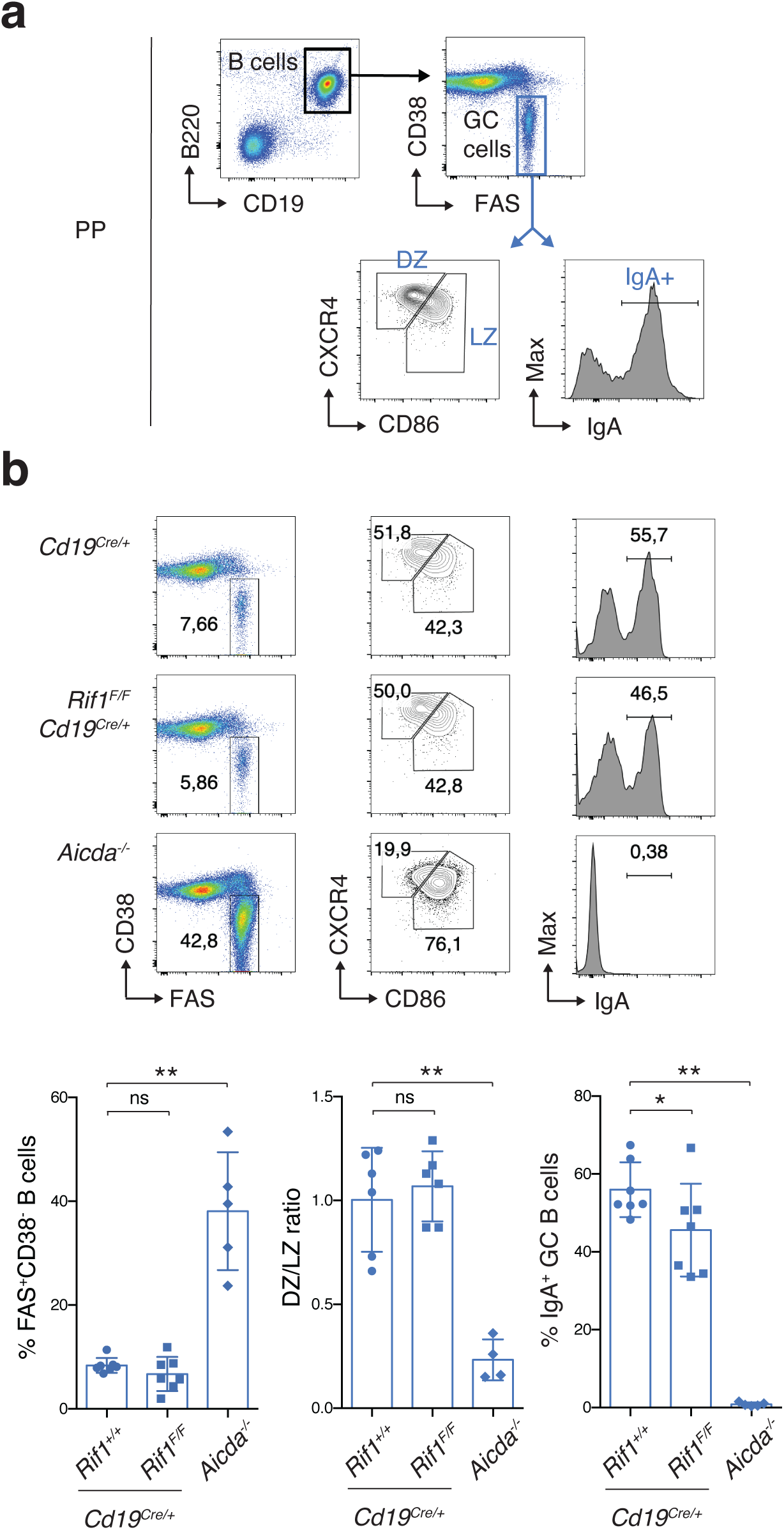
RIF1 deficiency does not affect GC B cell dynamics. **a** Gating strategy for Germinal Center (GC) analysis in Payer’s Patches (PPs). DZ: dark zone; LZ: light zone. **b** Top: Representative flow cytometry plots measuring percentage of GC B cells, DZ/LZ and IgA+ fraction of GC B cells in PPs of *Cd19^Cre/+^*, *Rif1^F/F^Cd19^Cre/+^ and Aicda^-/-^* mice. Bottom: Summary graphs for at least four mice per genotype in at least three independent experiments. Significance in panel **b** was calculated with the Mann–Whitney U test and error bars represent SD. ns: not significant; * = p ≤ 0.05; ** = p ≤ 0.01.

Interestingly, despite the severe CSR defect caused by RIF1 deficiency ^17–19,31^, PPs of *Rif1^F/F^Cd19^Cre/+^*mice displayed near-physiological levels of IgA^+^ GC B cells (Fig. 3, a and b), thus indicating that the few cells that successfully class-switch are actively selected forward in this context. We concluded that under conditions of chronic stimulation, RIF1 is dispensable for GC B cell formation, selection and proper DZ/LZ compartmentalization.

### RIF1 curtails plasma cell formation following immunization

Next, we analyzed the GC and PC compartment of control and *Rif1^F/F^Cd19^Cre/+^*mice immunized with the T-cell-dependent antigen 4-Hydroxy-3-nitrophenylacetyl hapten conjugated to Chicken Gamma Globulin (NP-CGG) (Fig. 4a). In contrast to chronic stimulation, NP-CGG immunization elicited a reduced number of GC B cells in the absence of RIF1 (Fig. S2, a and b). This phenotype is likely due to higher levels of apoptosis induced by unrepaired DSBs, as we have previously shown ^18^, or DNA end resection-dependent BCR loss ^41^. As expected for an acute immune response context, the levels of class-switched IgG1^+^ GC B cells (CD19^+^B220^+^FAS^high^CD38^low^IgG1^+^) were decreased in *Rif1^F/F^Cd19^Cre^* mice (Fig. S2, a and b).

**Figure 4.**
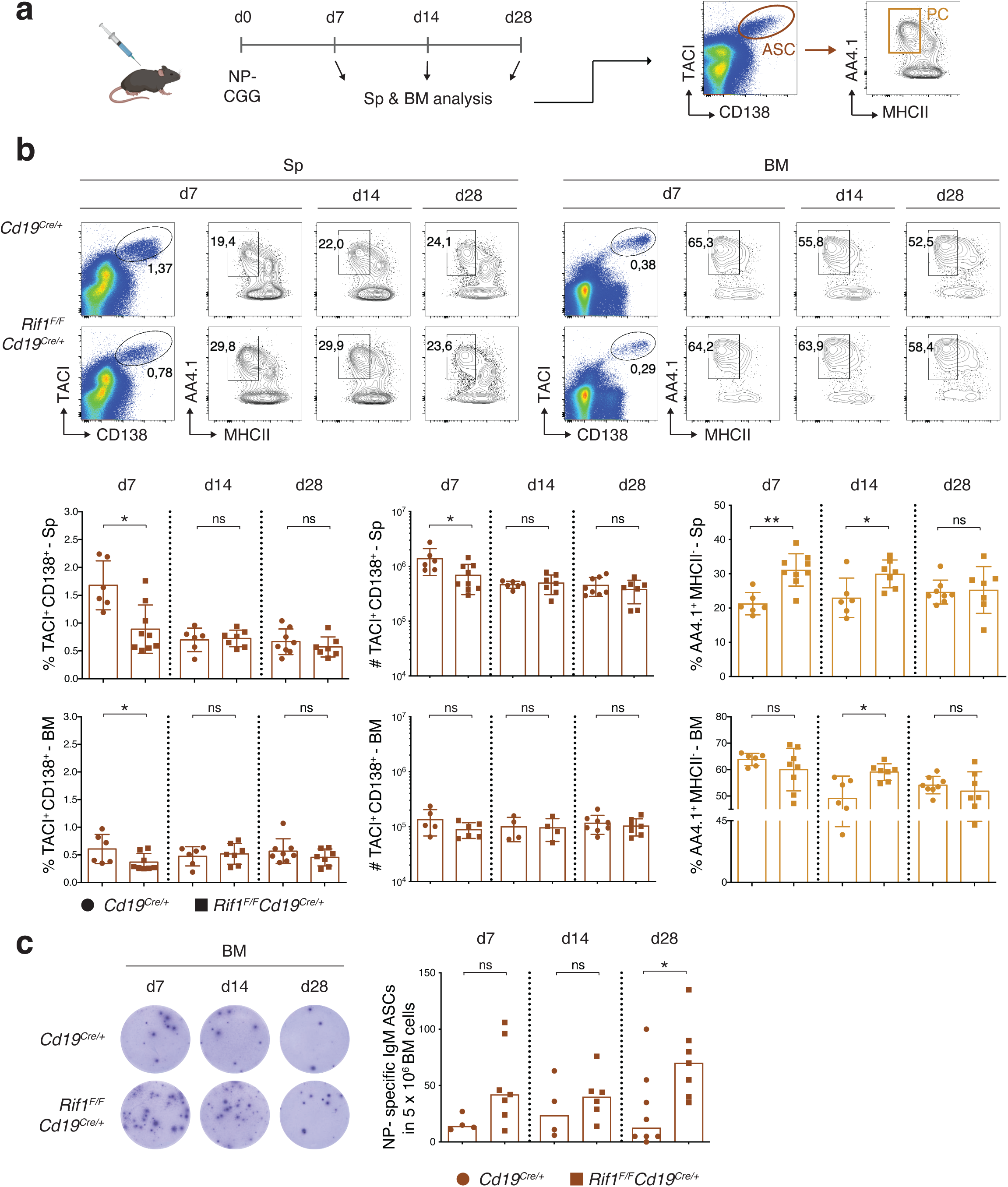
Ablation of RIF1 enhances plasma cell formation after immunization. **a** Schematic representation of the NP-CGG immunization protocol and gating strategy employed for the phenotypic analysis of the plasma cell compartment. d: day; NP-CGG: 4-hydroxy-3-nitrophenylacetyl hapten conjugated to Chicken Gamma Globulin; Sp: spleen; BM: bone marrow; ASC: antibody secreting cell; PC: plasma cell. **b** Top: Representative flow cytometry plots measuring percentage of antibody secreting cells and plasma cells in spleens and bone marrows of *Cd19^Cre/+^* and *Rif1^F/F^Cd19^Cre/+^* mice at the indicated days after immunization. Bottom: Summary graphs for at least six mice per genotype and time point in at least 3 independent experiments. **c** Left: Representative ELISpot analysis of NP-specific IgM ASCs in the BM at the indicated times after immunization. Right: Summary graph showing the number of NP-specific IgM ASCs per 5 x 10^6^ BM cells for at least four mice per genotype and time point. Significance in panels **b** and **c** was calculated with the Mann–Whitney U test. ns: not significant; * = p ≤ 0.05; ** = p ≤ 0.01. Error bars in panel **b** represent SD whereas the graph in panel **c** show the median of each dataset.

Despite the reduced number of GC B cells, *Rif1^F/F^Cd19^Cre^*mice displayed a consistent increase in the total pool of terminally differentiated PCs (TACI^+^CD138^+^AA4.1^+^MHCII^-^) compared to controls in both spleen and bone marrow at earlier time points post-immunization (day 7 and 14 in the spleen and day 14 in the bone marrow) (Fig. 4b). The higher proportion of PCs observed at day 7 in the spleen occurs on the background of a reduced ASC pool (Fig. 4b), which suggests that terminal B cell differentiation is accelerated in the absence of RIF1 also *in vivo*. The phenotype was no longer observed in either compartment at later time points (day 28, Fig. 4b) nor in unimmunized mice (Fig. S3 and ^17^). However, functional assessment of NP-specific PCs by ELISpot analysis showed a sustained trend towards higher levels of ASCs in the bone marrow of *Rif1^F/F^Cd19^Cre/+^* mice compared to controls, which was particularly pronounced at later time points after immunization (Fig. 4c). This latter observation is in agreement with the ability of terminally differentiated PCs to home to the bone marrow.

**Figure S3.**
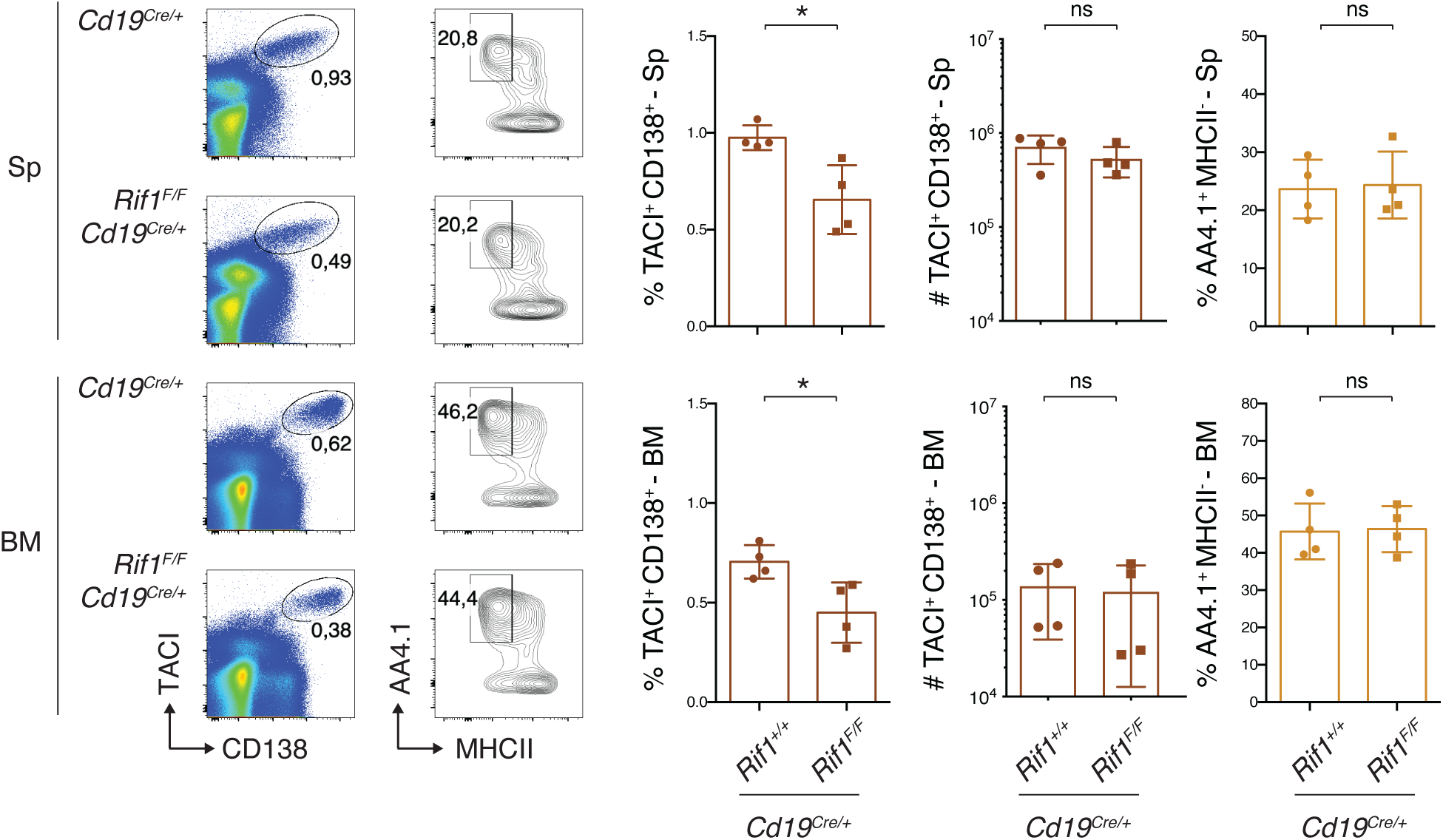
RIF1 ablation does not result in detectable changes of the plasma cell compartment in steady state condition. Left: Representative flow cytometry plots measuring percentage of antibody secreting cells and plasma cells in spleens and bone marrows of unimmunized mice of the indicated genotypes. The same gating strategy as in Figure 4a was employed. Sp: spleen; BM: bone marrow. Right: Summary graphs for four mice per genotype. Significance was calculated with the Mann–Whitney U test, and error bars represent SD. ns: not significant; * = p ≤ 0.05. Related to Figure 4.

**Figure S4.**
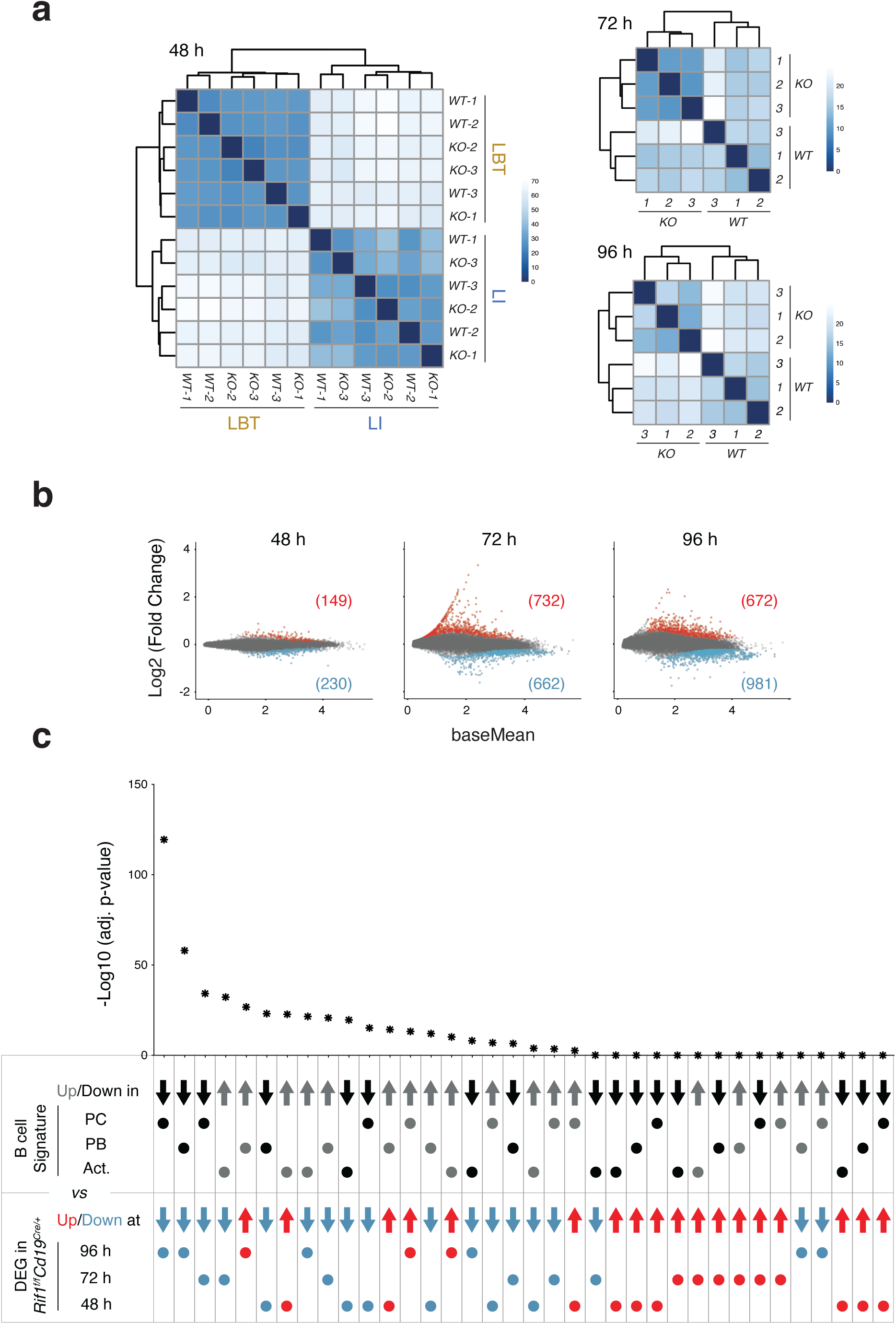
RIF1 deficiency alters the transcriptional landscape of activated B cells. **a-b** Dendrograms (**a**) and scatterplots (**b**) for the RNA-Seq datasets generated according to the experimental scheme depicted in Fig. 5a. For the 48 h time point, the analysis was performed on both LI- and LBT-stimulated cultures. Data summarizes results from three mice per genotype per stimulation condition. In the scatterplots, genes with an adjusted p-value <= 0.05 that are up- or down-regulated in *Rif1^F/F^Cd19^Cre/+^* cells are highlighted in red or blue, respectively, and the number of DEGs per category is included in parenthesis. **c** Statistical significance of pairwise comparisons between the indicated gene lists *via* hypergeometric analysis. Related to Figure 5.

These findings indicate that RIF1 deletion removes a physiological restraint imposed over the terminal differentiation of mature B cells *in vivo*, which, though concealed under steady-state conditions, is readily detectable at the systemic level upon immunization. Furthermore, taking into consideration also the *ex vivo* recapitulation of the increased ASC phenotype (Fig. 2) and unaffected GC B cell dynamics (Fig. 3), we concluded that RIF1’s ability to delay terminal B cell differentiation *in vivo* reflects a cell-intrinsic property independent from the GC reaction.

### RIF1 deficiency skews the transcriptional profile of activated B cells towards ASCs

RIF1 has been implicated in the modulation of transcriptional programs responsible for embryonic stem cell differentiation ^27,30,33,42^. Therefore, we asked whether the increased potential for terminal B cell differentiation in the absence of RIF1 reflects a transcriptional role in the modulation of B cell identity.

Activation of isolated splenic B cells with specific stimuli recapitulates several features of terminal B cell differentiation (Fig. 2 and ^10,43^). Hence, we monitored the consequences of RIF1 deficiency on the mature B cell transcriptome after *ex vivo* activation with LPS and IL-4. Comparative assessment of the transcriptional profiles of *Cd19^Cre/+^* and *Rif1^F/F^Cd19^Cre/+^*B cells identified only a limited number of considerably deregulated genes in the absence of RIF1 (N° genes with log2 FC < -1 and > 1 = 0, 105, and 47 at 48, 72, and 96 h, respectively) (Fig. 5, a and b, S4, a and b, and Table S1). Furthermore, the expression of the mature B cell identity transcriptional regulators (*Pax5*, *Ebf1*, *Foxo1,* and *Bach2*) was not affected (Fig. 5c). However, when we assessed the status of the key factors driving the ASC program (*Prdm1*, *Irf4*, and *Xbp1*), we found a near 2-fold increase in *Prdm1* transcript levels at 96 h after activation (Fig. 5, b and c). Since the *Prdm1*-encoded transcription factor BLIMP1 is required for the differentiation of pre-PBs into PBs and PCs ^8,10^, we cross-referenced the list of differentially expressed genes (DEG, adjusted p-value < 0.05) from *Rif1^F/F^Cd19^Cre/+^* activated B cells (Table S1) with the PB and PC signatures ^10^ (see Data analysis section of Methods). We observed a significant overlap between DEGs and the corresponding up-/down-regulated gene set in the ASC signatures, with the tendency being more pronounced for the down-regulated datasets (Fig. 5d, S4c and Table S2). We concluded that RIF1 deficiency in activated B cells results in a deregulated expression profile enriched in genes normally expressed in terminally differentiated B cells.

**Figure 5.**
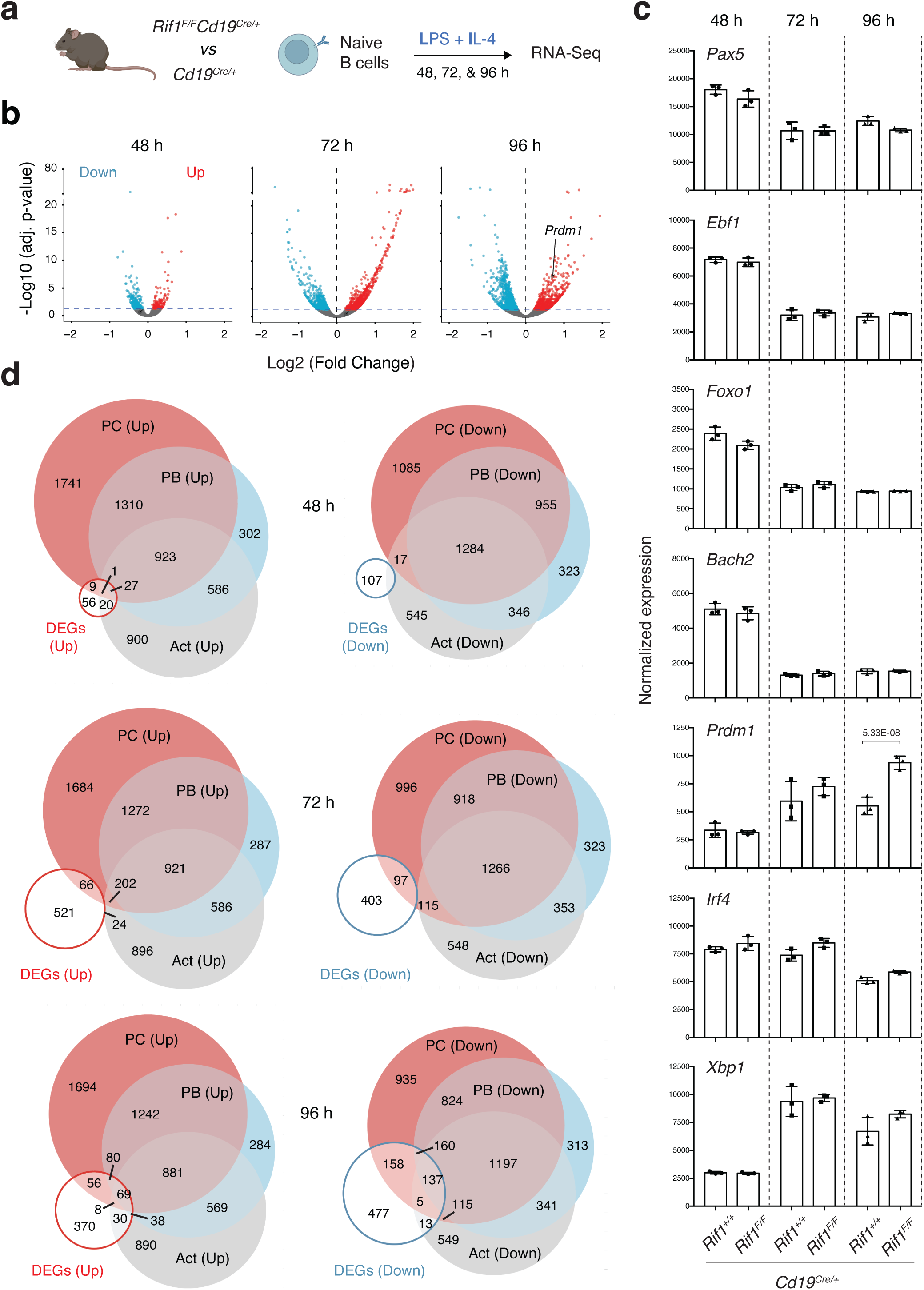
RIF1 deficiency skews the transcriptional profile of activated B cells towards ASCs. **a** Schematic representation of gene expression analysis in naïve B cells isolated from *Cd19^Cre/+^* and *Rif1^F/F^Cd19^Cre/+^* mouse spleens and stimulated *ex vivo* with LI for 48, 72, and 96 h. For the 48 h time point, the analysis was performed also on LBT-stimulated cultures. **b** Volcano plots displaying differentially expressed genes between control and RIF1-deficient splenocytes. The blue and red dots represent transcripts down- and up-regulated (adjusted p-value <= 0.05, blue horizontal dotted line), respectively, in *Rif1^F/F^Cd19^Cre/+^* cells. **c** Expression of *Pax5, Ebf1, Foxo1, Bach2, Irf4, Xbp1,* and *Prdm1* as determined by the RNA-Seq. Values were normalized by DESeq2 and the adjusted p-value of the only significant difference between samples is indicated. **d** Venn diagrams depicting the overlaps between genes up- (left) and down- (right) regulated in RIF1-deficient splenocytes at the indicated times post-activation and the corresponding up-and down-regulated (over naïve B cells) categories in the activated (Act) B cell, PB, and PC transcriptional signatures (Minnich et al. 2016).

### RIF1 binds genes involved in the modulation of adaptive immune responses

To uncover the molecular mechanism underlying RIF1 ability to maintain the transcriptional identity of mature B cells after activation, we monitored the genomewide occupancy of RIF1 in activated B cells from *Rif1^FH/FH^*mice under the same stimulation conditions employed for the *ex vivo* transcriptional analyses (Fig. 6a, ^25^). *Rif1^FH/FH^* splenocytes express physiological levels of a knock-in 1×<u>F</u>lag-2×<u>H</u>emagglutinin-tagged version of RIF1 (RIF1^FH^, ^44^) that supports its roles in mouse embryonic fibroblasts, embryonic stem cells, and B cells ^18,23,44^. We found that the vast majority of RIF1 peaks colocalized with promoters (56,4 %) and distal intergenic regions (25,4 %) (Fig. 6b). We next performed a functional enrichment analysis *via* Genomic Regions Enrichment of Annotations Tool (GREAT) ^45^ to assess the functional significance of both proximal- and distal-to-gene RIF1-binding events. We identified several categories of genes associated with the regulation of lymphocyte activation, function, and differentiation (Fig. 6c and Table S3). Altogether, these findings indicate that in activated B cells, RIF1 associates with cis-regulatory elements of genes involved in the modulation of the adaptive immune response.

**Figure 6.**
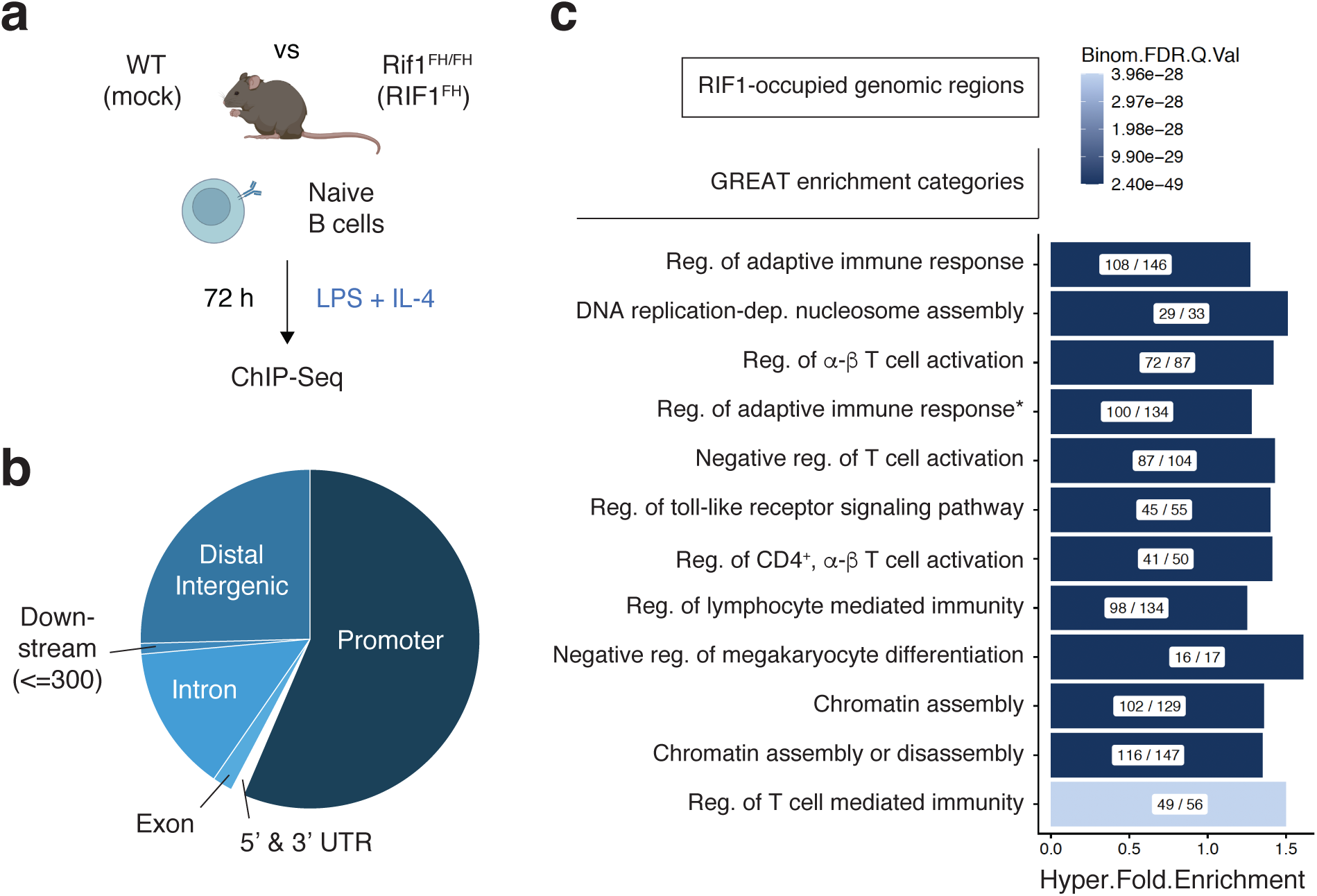
RIF1-occupied genomic regions in activated B cells comprise genes involved in lymphocyte activation and differentiation. **a** Schematic representation of the ChIP-Seq analysis in mature B cells isolated from spleens of *Rif1^FH/FH^* mice and stimulated *ex vivo* with LPS and IL-4 (LI). **b** Genomic distribution of RIF1-occupied annotated regions. **c** Gene ontology enrichment analysis as determined by GREAT for the genomic regions bound by RIF in LI-stimulated B cells. The complete name of the category marked with ” * ” is “Regulation of adaptive immune response based on somatic recombination of immune receptors built from immunoglobulin superfamily domain”. The X / Y ratio within each bar indicates the number of genes occupied by RIF1 (X) out of the total number of genes in the category (Y). Reg: regulation; FDR: false discovery rate.

### RIF1 counteracts the premature repression of BLIMP1 target genes

To further dissect how RIF1 modulates the transcriptional program driving B cell differentiation into ASCs, we next explored the relationship between RIF1 and BLIMP1, the key transcriptional regulator of terminal B cell differentiation ^10,25^. To this end, we first assessed the two factors’ relative genome occupancy by comparing RIF1 peaks (Fig. 6b) with BLIMP1-bound regions in activated B cells (see Methods). We found that 1300 genomic regions (corresponding to 1144 genes) were co-occupied by the two factors (Fig. 7a) and they primarily comprised active genes (Fig. 7b).

**Figure 7.**
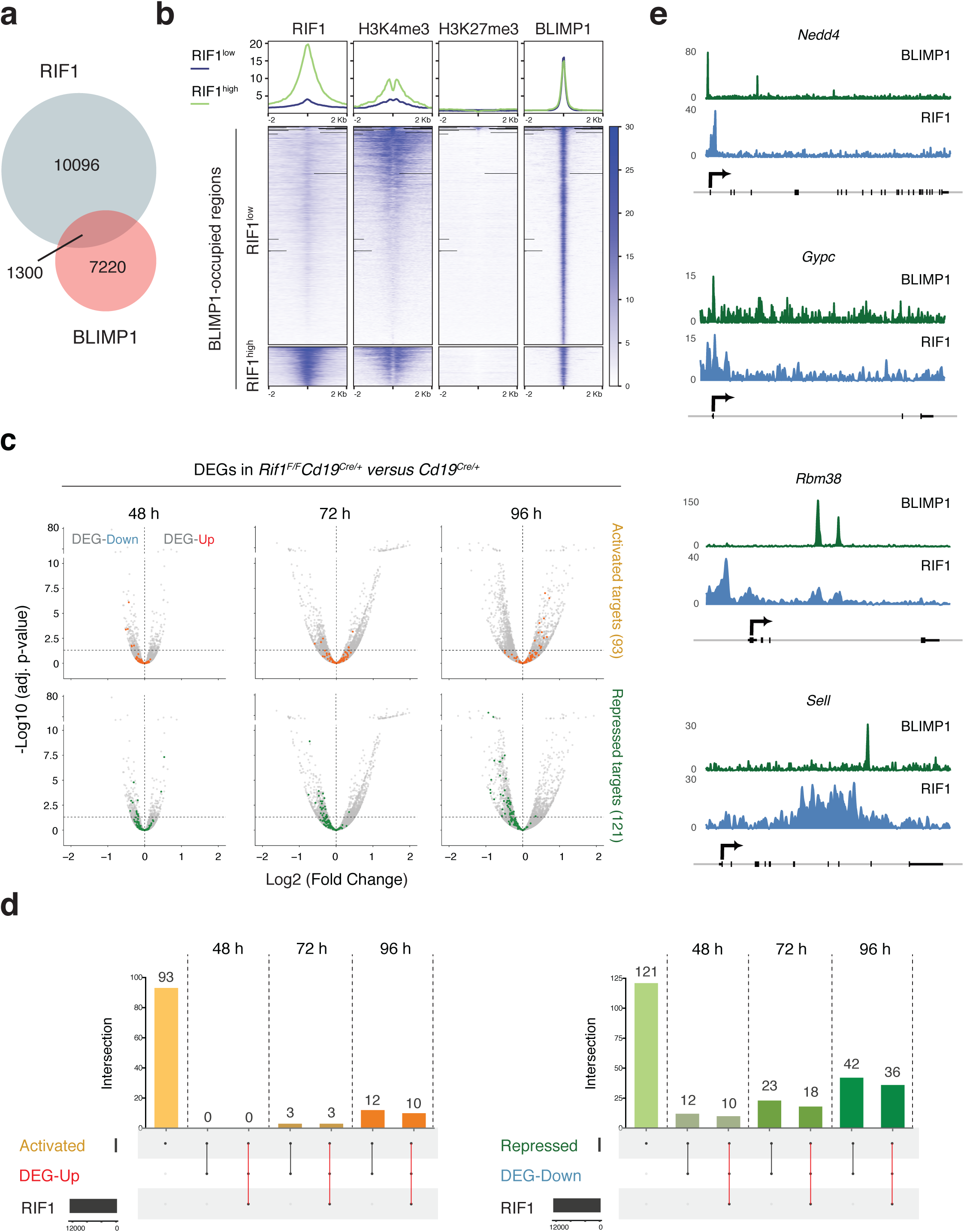
RIF1 counteracts the premature repression of BLIMP1 target genes. **a** Venn diagram depicting the overlap between RIF1- and BLIMP1-bound genomic regions in activated B cells. **b** Line plots (top) and heatmaps (bottom) depicting the comparative genome distribution of RIF1, H3K4me3, and H3K27me3 in reference to BLIMP1-occupied regions. The line plots illustrate the mean normalized signal distribution of the indicated proteins, whereas the heatmaps visualize their signal strength at each identified peak. The x-axis in both graph types represents the genomic region relative to the peak center. **c** Volcano plots displaying BLIMP1-activated (in orange, top) and -repressed (in green, bottom) target genes among the transcripts (in grey both up- and down-regulated) identified in the *Cd19^Cre/+^ versus Rif1^F/F^Cd19^Cre/+^* transcriptome analysis shown in Fig. 5. The horizontal dotted line in the plots denotes the adjusted p-value of 0.05. **d** UpSet plots depicting the intersection between activated BLIMP1 targets (Activated), genes differentially upregulated in *Rif1^F/F^Cd19^Cre/+^* B cells (DEG-Up) and RIF1-occupied regions (top), and between repressed BLIMP1 targets (Repressed), genes differentially downregulated in *Rif1^F/F^Cd19^Cre/+^* B cells (DEG-Down) and RIF1-occupied regions (bottom). The dot matrixes at the bottom of each graph show the intersection relationships among the data sets, with the number of common elements in the intersecting sets indicated above each bar. The red lines in the matrixes highlight the group of genes occupied and regulated by both BLIMP1 and RIF1. **e** ChIP-seq tracks for BLIMP1 (Minnich et al, 2016) and RIF1 at representative co-occupied BLIMP1 repressed targets from panel d.

Since gene binding is not indicative *per se* of regulatory activity, we next investigated the link between gene occupancy and transcriptional output. To this end, we first asked whether RIF1 deficiency affects the transcriptional status of BLIMP1 targets, which are defined as BLIMP1-occupied genes that are either up- (repressed, 121 targets) or down- (activated, 93 targets) regulated following its ablation ^10^. We found that several BLIMP1-activated genes were significantly up-regulated in *Rif1^F/F^Cd19^Cre/+^* cells at 96 h post-activation (Fig. 7c), which is line with the increased differentiation potential of these cultures (Fig. 5). More interestingly, BLIMP1-repressed targets were tendentially downregulated in the absence of RIF1, with the phenotype being evident from earlier time points after activation (Fig. 7c).

Next, we assessed RIF1 occupancy at BLIMP1 targets that are differentially expressed in the absence of RIF1. We found that a considerable portion of BLIMP1 repressed targets that are prematurely downregulated in the absence of RIF1 was also occupied by RIF1 in wild-type (*Rif1^FH/FH^*) cells at each time point post-activation, which is indicative of a direct RIF1 modulatory function on these genes (Fig. 7, d and e, and Table S4). This subset of targets comprises several genes whose repression has already been phenotypically and/or functionally linked to late B cell differentiation, including *Nedd4*, *Cd22*, *Ccr7*, *Btg1*, *Sell*, *B3gnt5*, *Bank1*, *Notch2*, and *Id3* (Fig. 7d and Table S4) ^10,11,46–55^. In contrast, the corresponding overlap for the upregulated BLIMP1 target genes was much more limited and only evident at later time points (Fig. 7d and Table S4). Altogether, these findings indicate that RIF1 supports the expression of several genes that are physiologically down-regulated by BLIMP1 as part of the transcriptional program promoting B cell differentiation into ASCs.

## Discussion

RIF1 has since long been shown to play a crucial role in humoral immunity because of its ability to protect AID-induced DSBs at *Igh*, thus ensuring productive class switching events ^17–19^. Here, we describe a unexpected modulatory role of RIF1 in regulating the timing of B cell differentiation to PCs.

Several recent studies have proposed that, besides SHM, BCR diversification *via* CSR can also influence the differentiation outcome of GC B cells as well as the PC transcriptome ^56–61^. Given the strict dependency of CSR on RIF1^17–19^, it is tempting to speculate that the skewed differentiation potential of RIF1-deficient B cells might represent an indirect consequence of defective isotype switching. However, the *ex vivo* recapitulation of the differentiation phenotype in the absence of RIF1 but not of its downstream DNA end protection partner SHLD1 or of AID (Fig. 2), argues against this possibility and in favor of a cell-intrinsic, DNA repair-independent role of RIF1 in fine-tuning PC differentiation kinetics.

Our results support a model where RIF1 serves as a regulator of terminal B cell differentiation through its capacity to counteract the premature repression of BLIMP1 target genes following activation. BLIMP1 is the master regulator of PC differentiation ^8,10,62^. Given that upregulation of BLIMP1 expression and appearance of PCs are closely intertwined events, it might appear challenging to unambiguously dissect the direct *versus* indirect aspects of RIF1 modulatory activity. However, the substantial overlap of RIF1 occupancy at BLIMP1 targets that are also differentially regulated in the absence of RIF1 is primarily observed at a subset of BLIMP1 repressed genes (Fig. 7). This finding indicates that RIF1 exerts its modulatory role on PC differentiation by directly controlling the transcriptional status of these loci rather than by indirectly influencing BLIMP1 expression. Mechanistically, the dynamic regulation of RIF1 expression in mature B cells supports this gene interaction co-regulatory model. We propose that high RIF1 levels prolong an active B cell status to enable efficient *Igh* DSB repair and CSR whereas its down-regulation contributes to a permissive environment for ASC differentiation (Fig. 1). Hence, our study has uncovered the existence of a fine-tuning regulatory mechanism that differentially impinges on the BLIMP1 transcriptional program controlling late B cell differentiation.

Previous reports have highlighted RIF1’s role in the differentiation of embryonic stem cells ^26,29,30,33^. In this context, RIF1 interacts with multiple Polycomb-group (PcG) proteins, and alters chromatin accessibility and target gene expression ^30,32,33^. Additionally, competition between RIF1 and KAP1 (and the long non-coding RNA *Tsix*) for interaction with *Xist* promoters establishes the asymmetric X-chromosome state at the onset of embryonic stem cell differentiation ^32^. A similar gene binding competition might apply to some of RIF1- and BLIMP1-co-occupied targets in the context of activated B cells, suggesting a potentially widespread molecular mechanism for RIF1-dependent transcriptional modulation across different cell types and differentiation states.

The multifaceted role of RIF1 underscores the evolutionary ingenuity in repurposing a single protein for diverse, yet equally critical, biological functions. RIF1 moonlighting in mature B cells has important implications for the pathological consequences of deregulated plasma cell generation. Whilst producing a diverse Igswitched repertoire enables effective antibody-mediated responses, tight regulation and fine tuning of late B cell differentiation is essential to counteract the development of autoimmunity and PC-derived malignancies ^63–65^. Hence, by enabling *Igh* diversification *via* CSR while exerting a modulatory function on PC differentiation, RIF1 integrates key requirements for the establishment of protective humoral immunity.

## Methods

### Mice and derived primary cell cultures

*Rif1^FH/FH^* ^44^, *Cd19^Cre^* ^66^, *Rif1^F/F^Cd19^Cre/+^* ^18^, *Shld1^-/-^* ^67^ and *Aicda^-/-^* ^38^ mice were previously described and maintained on a C57BL/6 background. Mice were kept in a specific pathogen-free (SPF) barrier facility and all experiments were performed in compliance with the European Union (EU) directive 2010/63/EU, and in agreement with Landesamt für Gesundheit und Soziales directives (LAGeSo, Berlin, Germany). Mice of both genders were used for the experiments.

Resting B lymphocytes were isolated from mouse spleens using anti-CD43 MicroBeads (Miltenyi Biotec), and grown in RPMI 1640 medium (Life Technologies) supplemented with 10% fetal bovine serum (FBS, Sigma-Aldrich), 10 mM HEPES (Life Technologies), 1 mM Sodium Pyruvate (Life Technologies), 1X Antibiotic Antimycotic (Life Technologies), 2 mM L-Glutamine (Life Technologies), and 1X 2- Mercaptoethanol (Life Technologies) at 37 °C and 5% CO2 levels. Naïve B cells were activated by addition of 5-25 μg/ml LPS (Sigma-Aldrich) and 5 ng/ml of mouse recombinant IL-4 (Sigma-Aldrich) (L-I), or 5 μg/ml LPS, 10 ng/ml BAFF (PeproTech) and 2 ng/ml TGFβ (L-B-T), or 5 μg/ml LPS only (L).

### Western Blot

WB analysis of protein levels was performed on whole-cell lysates prepared by lysis in radioimmunoprecipitation assay buffer (Sigma-Aldrich) supplemented with Benzonase (Sigma-Aldrich), Complete EDTA free proteinase inhibitors (Roche) and Pierce Phosphatase Inhibitor Mini Tablets (Thermo Fisher Scientific). The antibodies used for WB analysis are anti-RIF1 ^18^ and anti-β-actin (Sigma).

### RNA-Seq

For each RNA-Seq dataset, the analysis was performed on three mice per genotype. Splenocytes were cultured in LPS and IL-4 (LI), LPS, BAFF and TGFβ (LBT), or LPS (L), and cells were collected at the indicated time points by centrifugation. RNA was extracted with TRIzol (Invitrogen) according to manufacturer’s instructions, and ribosomal RNA was depleted using Ribo-Zero Gold rRNA Removal Kit (Illumina) for all datasets except for the RNA-Seq analysis in naïve versus LI/LBT/L-activated splenocytes from WT mice (Fig. 1 and S1), for which RNase H (Epicentre) treatment was used. Libraries were prepared with TruSeq Stranded Total RNA Library Prep Kit Gold (Illumina), and run in one lane on a flow cell of NovaSeq 6000 SP (Illumina).

### ChIP-Seq

ChIP-Seq for RIF1 in LPS and IL-4-stimulated splenocyte cultures was previously described ^25^. H3K4me3 ChIP-Seq was performed in splenocytes activated with LPS and IL-4 for 72 h, and employed anti-H3K4me3 antibody (abcam, ab8580) for the ChIP part of the previously described protocol ^68^. For H3K27me3 ChIP-Seq, we used H3K27me3 antibody (Cell Signalling, C36B11), and 2.5% human shared chromatin was spiked into all samples as an internal reference for normalization ^69^.

### B cell proliferation, development and differentiation analyses

For analysis of CSR in *ex vivo* cultures, cell suspensions were stained with anti-IgG1-APC (BD Biosciences), anti-IgG3-Biotin and Streptavidin-APC (BD Biosciences), or anti-IgG2b-PE (BioLegend). For analysis of plasmablast differentiation *ex vivo*, isolated naïve splenic B cells were cultured at a density of 0,5 x 10^6^ cells/ml in the presence of 25 μg/ml LPS and 5 ng/ml IL-4 and stained with anti-CD138 (BioLegend). B cell proliferation was assessed by CellTrace Violet (Thermofisher) dilution according to manufacturer’s instructions. For analysis of plasma cell-like differentiation *ex vivo*, the induced GC B (iGB) culture system was used as described before ^35,36^. Briefly, 40LB feeder cells (Balb/c 3T3 cell line expressing exogenous CD40-ligand (CD40L) and B-cell activating factor (BAFF)) were irradiated with 80 Gy and co-cultured with primary splenic B cells for 4 days in high glucose DMEM medium (Gibco) supplemented with 10% FBS, 10 mM HEPES, 1 mM Sodium Pyruvate, 1X Pen/Strep (Life Technologies), 1x MEM Non-Essential Amino Acids (Life Technologies), 2 mM L-Glutamine, 1X 2-Mercaptoethanol and 1 ng/ml IL-4 at 37 °C and 5% CO_2_ levels. On day 4, cells were harvested, washed one time with PBS and re-plated in a newly irradiated 40LB feeder layer in the presence of 10 ng/ml IL-21 (PeproTech) for the next 4 days. For assessing CSR and plasma cell-like differentiation on day 4 and 8, 1 x 10^6^ B cells were washed one time with FACS buffer (PBS supplemented with 1% FCS and 1 mM EDTA), blocked with TruStain fcX (BioLegend) for 10 min at 4° C and stained with the respective antibodies as stated above. 1 µg/ml of propidium iodide (PI) was used for live/dead cell staining.

For analysis of germinal centers and plasma cell compartments *in vivo*, 8-14 week-old mice were sacrificed to isolate the spleen, tibia, and Payer’s patches (PP). Single cell suspensions from spleen and tibia were incubated for 2 min with ACK buffer (Gibco) for red blood cell lysis. For surface staining, 5-7 × 10^6^ cells were first blocked with TruStain fcX for 10 min at 4 °C and then stained for 20 min at 4° C. The antibodies employed for the staining were detecting: CD138, CD267/ TACI (BD Pharmingen), MHC-II (Biolegend) and CD93/AA4.1 (Biolegend) for the plasma cell compartment; CD19, B220, CD86 (Biolegend), CD184/CXCR4, CD95/FAS (BD Pharmingen), IgA and CD38 (eBioscience) for PP analysis; and CD19, B220, CD38, CD95/FAS and IgG1 (BD Bioscience) for splenic GC analysis. Cells were resuspended in FACS buffer containing PI and analyzed. Immunization was performed by intraperitoneal injection of 100 µg of NP-CGG (Biosearch Technologies; ratio 10-20) precipitated in alum (Sigma).

All samples were acquired on a LSRFortessa cell analyzer (BD Biosciences).

### ELISpot assay

To detect NP-specific plasma cells, MultiScreen_HTS_-IP plates (Merck) were activated with 35% ethanol in PBS, washed three times with PBS, and finally coated with 2 μg/mL NP-BSA (loading ratio >20, BioCat) at 4°C overnight. Next day, plates were washed twice with PBS and blocked with ELISpot medium (RPMI 1640 supplemented with 10% FBS and 1% pen/strep) for 2 h at 37 °C. 5 x 10^6^ total spleen and BM cells (as well as serial five-fold dilutions) were added to the NP-BSA coated plates and cultured overnight in ELISpot medium. The following day, wells were washed six times with PBS supplemented with 0.1 % Tween20 and incubated for 2 h with 1 μg/mL biotinylated goat α-mouse IgM or IgG1 (SouthernBiotech) at 37 °C. Spot visualization was performed by incubation with 0.3 U/ml streptavidin-AP conjugate (Roche) for 30 min at room temperature followed by three washes with running distilled water, equilibration in AP buffer (100 mM Tris-HCl, pH 9.0, 150 mM NaCl, and 1 mM MgCl_2_), and development using NBT/BCIP substrate mix (Promega) diluted in AP buffer. Plates were scanned and spots were counted using the ImmunoSpot® Series 6 Alfa Analyzer and ImmunoCapture™ Image Acquisition as well as ImmunoSpot® Analysis software (C.T.L.).

### Data availability

All RNA-Seq datasets reported in this study, and H3K4me3 and H3K27me3 ChIP-Seq data have been deposited in the GEO repository under accession number GSE237560. The transcriptional signatures of activated B cells, PBs, and PCs have been defined using the corresponding (and naïve B cells for baseline comparison) RNA-Seq datasets from (^10^, GSE71698). RIF1 ChIP-Seq was previously reported (^25^, GSE228880). The BLIMP1-bound regions in activated B cells were defined based on the BLIMP1 Bio-ID dataset from (^10^, GSE71698). The lists of BLIMP1-activated and - repressed targets have been previously described ^10^.

### Data analysis and Softwares

#### RNA-Seq

All RNA-Seq datasets, whether specifically generated for this study or obtained from public databases, have been analyzed/re-analyzed using the same pipeline implemented for this project. The RNA-seq data was analyzed using the Galaxy platform ^70^. HTSeq ^71^ mapped the reads to the GRCm38 assembly for mouse and GENCODE-M25 (www.gencodegenes.org) was used to count the transcript abundance. Differential expression analysis was done *via* the DESeq2 ^72^ package for R, which uses the Wald test for significance. To define the transcriptional signatures of activated B cells, PBs, and PCs, the analysis was performed using the naïve B cell RNA-Seq dataset from the same study ^10^ for baseline comparison. Expression of differentially expressed genes were obtained from Immgen database ^34^ and visualize by R and ggolot2 packages (https://ggplot2.tidyverse.org/) after scaling. For the comparative assessment of *Cd19^Cre/+^* and *Rif1^F/F^Cd19^Cre/+^*B cell transcriptomes at 48 h after activation, RNA-seq datasets from L-I and L-B-T splenocytes cultures were merged. The increase in sample size (from 3 to 6 repeats) for both groups enhanced statistical power to detect minor gene expression differences. The multiple treatments are used to discern genotype effects, thus ensuring that the observed differences are due to the absence of RIF1.

#### ChIP-Seq

ChIP-Seq and BLIMP1 Bio-ID datasets have been analyzed/re-analyzed using the same pipeline implemented for this project. FASTQ files were aligned against mouse genome (mm10) using BWA aligner ^73^. Processing and peak-calling of ChIP-Seq data were performed with MACS2 ^74^. Peak annotation was done using R and ChIPseeker package ^75^. Functional assessment of genomic regions enrichment and motif analysis were performed by GREAT and MEME-ChIP, respectively ^45,76^. For the comparative genomic distribution analysis of RIF1, H3K4me3, and H3K27me3 in reference to BLIMP1-occupied regions, the aligned reads were converted into BigWig format. The signals were subsequently transformed into a matrix *via* the ComputeMatrix software, and visualized as a heatmap^77^.

#### Statistical analysis

The statistical significance of differences between groups was determined by the Mann–Whitney U test for all data presented in this study, with the following exceptions. For the RNA-Seq analysis, the expression values of specific transcripts were normalized by DESeq2, and genes with an adjusted p-value < 0.05 were considered to be significantly differentially expressed. Statistical details of experiments can be also found in the figure legends.

#### Hypergeometric analysis

Gene expression data were sourced from experiments from this study and published datasets^10^ (GSE71698) and compiled into a single CSV in parallel to a universe of genes (list of all known genes in mouse genome, mm10). A custom R function was developed to conduct overrepresentation analysis *via* hypergeometric testing. This function computes the likelihood of observing the intersection of genes between two lists, considering the total gene count in the universe and adjusting for the sizes of the lists involved. Pairwise comparisons between all unique pairs of gene lists were conducted by the hypergeometric test, which was implemented using the phyper function from R’s base stats package, with the test’s directionality set to identify overrepresented genes. The resulting p-values were adjusted for multiple comparisons using the Bonferroni correction method to control for the family-wise error rate. For a comprehensive overview of the pairwise comparison results, we reported the Bonferroni-adjusted p-values as -log10 values. This transformation enhances the interpretability of the results, with higher values indicating more significant overlaps between gene list pairs.

#### Source Codes for Data Analysis

All codes used to generate graphs and to compare different datasets can be found on our GitHub page (https://github.com/arahjou/RIF1_paper.git).

## Acknowledgments

We thank all members of the Di Virgilio lab for discussion; L. Keller and T. Rüster (Di Virgilio lab, MDC, Berlin) for genotyping; W. Winkler and C. Farre i Diaz (Janz/Mathas and K. Rajewsky labs, MDC) for *in vivo* and ELISpot protocols; and D. Pasini (IEO, Milan) and G. Gargiulo (MDC) for their valuable feedback. Figure schematics were created using images from BioRender. The project was funded by the Helmholtz-Gemeinschaft Zukunftsthema “Immunology and Inflammation” ZT-0027 (to M.D.V.) and the Initiative and Networking Fund of the Helmholtz Association (to M.D.V.).

## Author contributions

A.R. and E.K. conceived the project idea, designed and performed experiments; M.B.L. and V.D.B prepared the RNA-seq libraries; A.R. analyzed all high-throughput sequencing data; R.A. contributed to the sequencing data analysis; R.P. engaged in active discussions on the study; M.D.V. secured the funding for the project, supervised all aspects of the study, and wrote the manuscript; A.R. and E.K. reviewed and edited the manuscript.

